# Associative nitrogen fixation could be common in South African mesic grassland

**DOI:** 10.1101/2022.07.07.499153

**Authors:** Craig D Morris, Danvir R Ramesar, Richard J Burgdorf

## Abstract

Non-symbiotic nitrogen-fixing bacterial diazotrophs closely associated with the roots of grasses probably contribute most of the new nitrogen acquired to sustain productive natural grasslands, yet their ecology is poorly understood, especially in southern Africa. We looked for genetic evidence, using qPCR and gel electrophoresis, for the presence of the bacterial *nifH* gene associated with the roots of four grass species (20 plants each) in a mesic grassland in South Arica, which would indicate the potential for N fixation by diazotrophs. Grasses most tolerant of low N (*Aristida junciformis*) were predicted to harbour the most diazotrophs, especially compared to those most responsive to fertiliser N (*Eragrostis curvula*). However, the *nifH* gene was found in all 80 root samples and did not differ in copy number between species. Sequencing of a representative sample confirmed the identity of the *nifH* gene. The recently burned half of the grassland had 60% more *nifH* genes than the area burned 15 months previously, suggesting that grass growth stimulated by fire could recruit diazotrophs. Given their ubiquity and importance in the N economy of grasslands, research is required to characterise root-associated diazotroph communities, quantify their N fixation rates, and understand their environmental controls.

## Introduction

Nitrogen (N) is an essential metabolic building block for plant production and the key nutritional currency in grassland herbivory food webs, including in domestic livestock production systems (Woodmansee et al. 1981; Mason et al. 2022). Biological nitrogen fixation (BNF) by microorganisms is the major source of N input to natural grassland ecosystems (Kleiner 1975; Cleveland et al. 1999; Fowler et al. 2013). Microorganisms are also the chief recyclers of extant organic N in soil, converting N into inorganic compounds (ammonia and nitrate) available for uptake by plants (Paul and Juma 1981; Risch et al. 2019). Herbaceous and woody legumes that form symbiotic relations with root-nodulating nitrogen-fixing bacteria (diazotrophs) are generally assumed to be the main means of BNF in temperate (Becker and Crockett 1976) and tropical grassland (Pellegrini et al. 2016; Reis et al. 2020). There is, however, growing evidence (since the 1970s) that non-symbiotic (free-living) soil diazotrophs with significant nitrogenase activity could associate with the roots of several grass species, providing them with plant-available N (e.g., Nelson et al. 1976; De-Polli et al. 1977; Boddey and Knowles 1987).

A notable example of associative nitrogen fixation (ANF) in grasses is the Brazilian varieties of sugarcane that sustain high yields with little mineral fertiliser input because of N supplied by a variety of leaf and stem endophytes and rhizosphere/rhizoplane diazotrophs (Boddey et al. 2003). Estimates of the amount of plant N needs supplied by ANF to cultivated bioenergy and forage grasses range, for example, from <20% for *Miscanthus* × *giganteus* (Keymer and Kent 2014), 24–38% for *Panicum maximum* (Miranda and Boddey 1987), to 18–70% for *Pennisetum purpureum* (de Morais et al 2012). Inputs from ANF are important for maintaining perennial grasses on poor soils (Gupta et al. 2019). However, measuring *in situ* rates of ANF in multi-species natural grasslands and estimating the amount of fixed N absorbed and incorporated in plant tissues is challenging (Khan et al. 2021), especially when conditions favouring N-fixation can be episodic (Roley et al. 2019).

Conditions for optimum ANF and the ecology of non-symbiotic diazotrophs in grassland ecosystems are currently poorly understood (Roley et al. 2019; Smercina et al. 2019), particularly for African tropical and subtropical grasslands. In the Serengeti, Ritchie and Raina (2016) detected the nitrogenase gene, *nifH*,closely associated with the roots of *Themeda triandra* and measured N_2_ uptake by root sections, which was positively correlated with *nifH* copy number at rates comparable to that of a native legume. They concluded that the large amounts of N fixed by root endophytes could contribute significantly to the nitrogen economy of that grassland. In South Africa, the ability of the *nifH* gene to reduce acetyl *in vitro* was used by Staphorst and Strijdom (1987) to indirectly identify and then culture diazotrophs associated with the roots of nine pasture and natural grassland (veld) grasses. They also noted that many other local grasses could most likely also be capable of ANF. Similarly, Maasdorp (1987) found the highest potential for grass ANF in wet areas in Zimbabwe, with *Paspalum urvuillei* estimated to fix up to 76 kg N ha^-1^ yr^-1^. Haiyambo et al. (2015) isolated 21 diazotrophic strains associated with five grasses in Namibia. Apart from these few studies, the occurrence of diazotrophs and the potential for biologically significant ANF in South African grasslands have been largely overlooked.

In this study, we looked for direct genetic evidence for diazotrophs associated with the roots of four grasses differing in sensitivity to N in a mesic grassland in KwaZulu-Natal, South Africa. We screened root samples for the presence of the *nifH* gene - a highly conserved, reliable marker of N_2_-fixation (Reed et al. 2011) - on or inside their roots. We expected *nifH* gene copy number would be highest on grasses dominant on dystrophic soils with low N where they could be relying on ANF to meet their N needs and lowest or absent from the roots of fast-growing grasses that are effective and efficient at acquiring N from soil (Marques et al. 2017).

## Materials and Methods

The four grasses used, in order of decreasing yield response to applied inorganic fertiliser N, were: *Eragrostis curvula, Themeda triandra, Tristachya leucothrix*, and *Aristida junciformis* (Morris 2016). *Eragrostis curvula* thrives in high-N plots, *A. junciformis* can dominate on very poor soils and the other two species are common in unfertilised mesic grassland, which is typically low in N (Fynn and O’Connor 2005; Morris 2016). Root samples (down to 150 mm) from 20 tufts of each species were collected from a diverse mesic remnant grassland (KwaZulu-Natal Hinterland Thornveld; Mucina and Rutherford 2006) in Pietermaritzburg, South Africa. Ten of these replicates were from one half of the ∼5 ha field that had been burned in late winter 15 months previously (hereafter referred to as the ‘old’ sward) and the other ten in a ‘young’ sward burned five months previously.

Roots were shaken vigorously for 10 min to remove surrounding soil and washed with distilled water (Barillot et al. 2013), leaving only potential diazotrophs inside roots and or adhered to their surface (i.e., the rhizoplane). Root samples were stored at -20°C in the laboratory. Frozen root samples were ground in liquid nitrogen using a mortar and pestle and DNA was extracted using a Macherey-Nagel NucleoSpin™ Plant II kit. All metagenomic sample DNA extracts were standardized to 10 ng ul^-1^ in a 1 x TE buffer.

As per Ritchie and Raina (2016), the *nifH g*ene was amplified using real-time or quantitative PCR (qPCR). A *nifH* gene qPCR standard was prepared from genomic DNA of a commercial *Rhyzobium* sp. culture. All qPCR reactions were prepared as 20 µL solutions that contained: 1 x KAPA SYBR Fast Universal Fast qPCR Master Mix (KAPA Biosystems, U.S.A.), 0.5 µM *nifH* forward primer (CSATCAACTTCCTBGARGA), 0.5 µM reverse primer (GCCATCATBTCRCCGGA) and 1 ng µL^-1^ of template DNA. All qPCR reactions were conducted on the QuantStudio 5 qPCR machine (Thermo Fisher Scientific, U.S.A.). The amplification parameters used included an initial denaturation of 95°C for 10 min followed by 40 cycles of 95°C for 5 s, 55 – 50°C for 20 s (stepped down by 0.5°C for the initial 10 cycles) and 72°C for 20 s. Terminal elongation was performed at 72°C for 3 min. All samples were processed in the same qPCR run, which included duplicate qPCR *nifH* standards (10^3^, 10^4^, 10^5^ and 10^6^ *nifH* copies) for copy number determination. All qPCR reaction products were resolved by electrophoresis on a 1.5% agarose gel at 8 V cm^-1^ for 60 min to confirm the presence of the ∼200 bp *nifH* gene fragment.

The positive *Rhizobium* control and eight randomly selected PCR amplicons were sent to Inqaba Biotechnical Industries (Pty) Ltd, South Africa for Sanger sequencing in forward and reverse directions. Consensus sequences were produced in BioEdit (version 7.2.5). Consensus sequences were trimmed to include regions within forward and reverse primers and were submitted to a Genbank nucleotide BLAST (Altschul et al. 1990) with models and environmental samples excluded from the search (blastn). The best matches were reported to confirm that the fragments that were sequence were of *nifH* gene origin. Control and sample sequences were compared to ensure that the *nifH* amplicon presence was not due to contamination or technical error.

Variability between species in *nifH* gene number was tested by one-way ANOVA (n = 80) whereas within-species differences between unreplicated locations (old vs new sward) were compared with a Student’s t-test. Gene copy number was square-root transformed to normalise and homogenise residual variance.

## Results

The qPCR and gel electrophoresis revealed the presence of suitably sized (∼200 bp) *nifH* gene target amplicons in all 80 root samples (Figure 1). The *nifH* gene copy number varied over 1000-fold, from 179 to 245,350 copies ng^-1^ template DNA.

**Figure 1:**
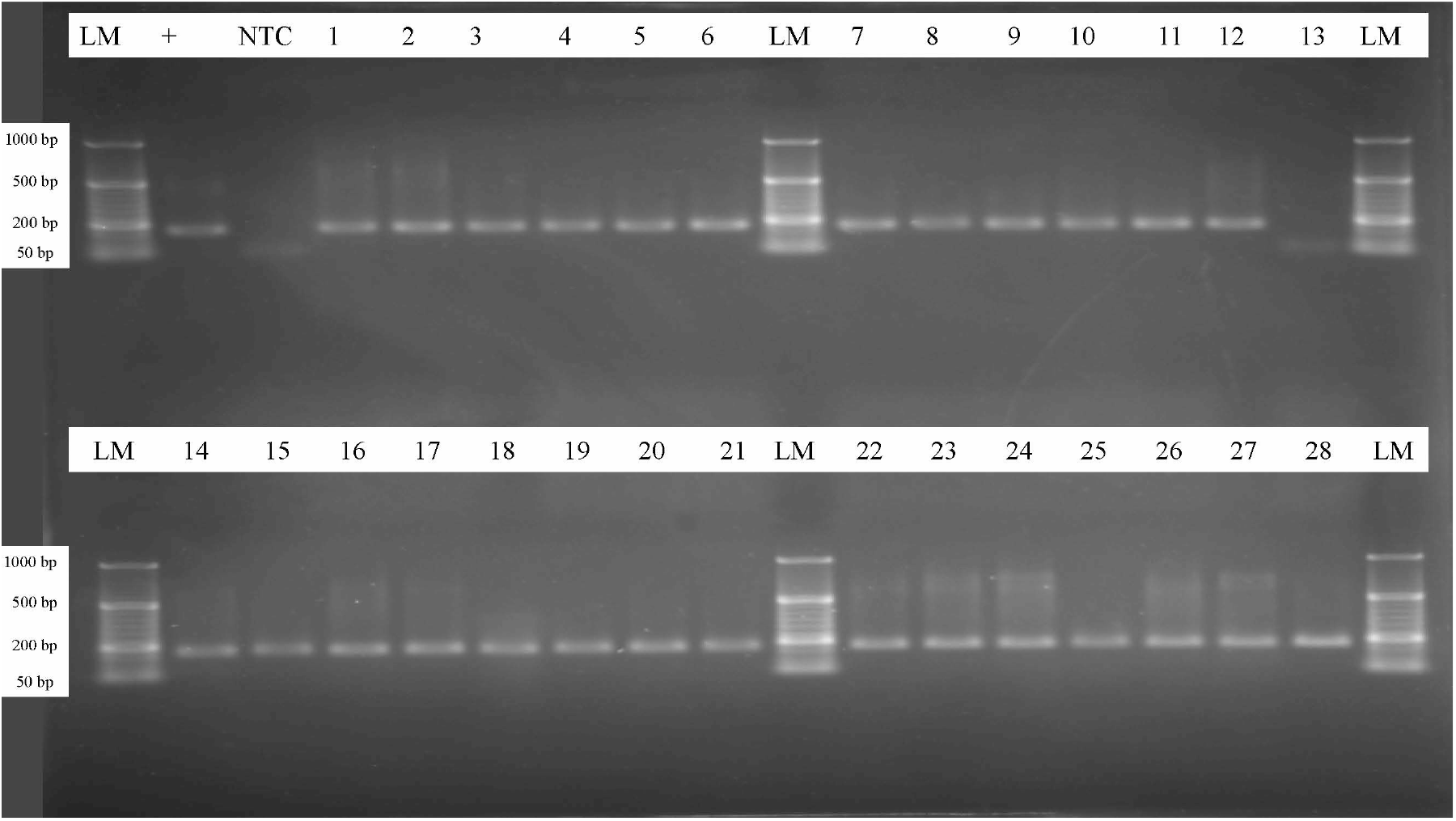
Example Agarose (1.5%) gel electrophoresis of *nifH* amplicons. Samples loaded included a 50 bp ladder marker (LM), *Rhizobium* sp. as the positive (+) control, a no template control (NTC). PCR amplicons of soil samples designated from 1 – 28 were: NTT R1, NTT R2, NTT R3, NTT R4, NTT R5, NTT R6, NTT R7, NTT R8, NTT R9, NTT R10, OTT R1, OTT R2, OTT R3, OTT R4, OTT R5, OTT R6, OTT R7, OTT R8, OTT R9, OTT R10, NAJ R1, NAJ R2, NAJ R3, NAJ R4, NAJ R5, NAJ R6, NAJ R7, NAJ R8 respectively. N = new sward, O = old sward. TT = *Themeda triandra*, AJ = *Aristida junciformis*. R1-10 = replication number.

The mean number of copies of the *nifH* gene did not vary significantly between grass species (F_3,76_ = 1.769, *p* = 0.160). The old and new swards did, however, differ systematically across all species in mean gene copy number (*t*_78_ = 3.375; *p* = 0.0012) with the mean copy number 60% higher (untransformed scale) for grass roots in the newly-burned than the old, mature sward (Figure 2).

**Figure 2:**
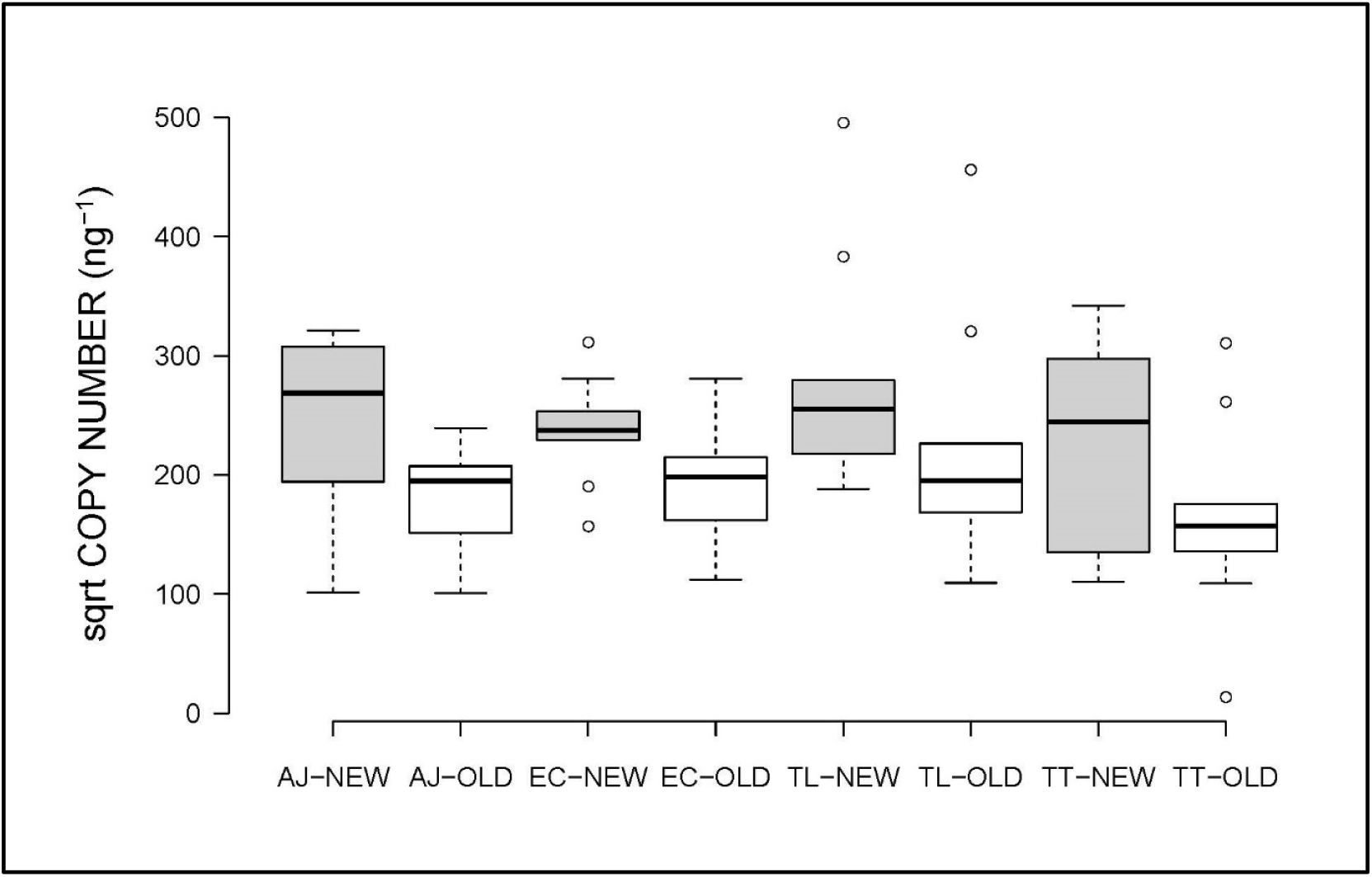
Boxplot of median, interquartile range, and outliers for *nifH* gene copy number associated with the roots of four grass species in a mesic grassland. AJ = *Aristida junciformis*, EC = *Eragrostis curvula*, TL = *Tristachya leucothrix*, TT = Themeda *triandra*. OLD = old sward (unfilled boxes), NEW = new sward (filled boxes).

Trimmed *nifH* gene sequences were approximately 150 bp in length. Sequences had high levels of ambiguities but the dominant *nifH* gene sequence in each sample had the closest similarity to the *nifH* gene sequences from *Bradyrhizobium* species (Table 1).

**Table 1.** Sequence matches for selected amplicons and NCBI database sequences associated with the roots of four mesic grasses in the Old and New field location

| Sequence No. & Sample ID | Matching sequence description | Coverage (%) | Identity (%) | Accession |
| --- | --- | --- | --- | --- |
| S1 - *Rhizobium* sp. control | *Bradyrhizobium* sp. *NifH* gene | 100 | 94.12 | KR491971.1 |
| S7 - New *T. triandra* R5 | *Bradyrhizobium* sp. chromosome | 99 | 92.16 | CP059398.1 |
| S32 - New *A. junciformis* R10 | *Bradyrhizobium* sp. *NifH* gene | 99 | 89.74 | MK559280.1 |
| S39 - Old *A. junciformis* R7 | *Bradyrhizobium* sp. *NifH* gene | 99 | 90.79 | MK559280.1 |
| S50 - New *E. curvula* R8 | *Bradyrhizobium* sp. *NifH* gene | 99 | 92.26 | KT965790.1 |
| S60 - Old *E. curvula* R8 | *Bradyrhizobium* sp. *NifH* gene | 99 | 91.5 | MK559280.1 |
| S62 - Old *E. curvula* R10 | *Bradyrhizobium* sp. *NifH* gene | 99 | 90.91 | MK559283.1 |
| S66 - New *T. leucothrix* R4 | *Bradyrhizobium* sp. *NifH* gene | 99 | 90.32 | MK559280.1 |
| S77 - Old *T. leucothrix* R5 | *Bradyrhizobium* sp. complete sequence | 99 | 87.26 | CU234118.1 |

## Discussion

Our finding that the *nifH* gene was ubiquitous, on or inside the roots of all four species across the study area supports the assertion that N_2_-fixing bacteria could be common associates of grasses in subtropical, mesic grassland in South Africa (Staphorst and Strijdom 1987). The presence of the gene coding for nitrogenase indicates the potential for non-symbiotic nitrogen fixation by subtropical grasses. Rates of nitrogen fixation by local grasses have not been measured but ANF could provide a critical portion of the nitrogen growth needs of grasses (Keuter et al. 2014; Smercina et al. 2019). This ‘new’ N could contribute substantially to the nitrogen economy of mesic grassland (Ritchie and Raina 2016), probably far more in total than that fixed by indigenous herbaceous legumes (Reed et al. 2011) which are far less abundant than grasses in South Africa (Trytsman et al. 2019; Vázquez et al. 2022). Such ANF could also be one of the reasons, along with microbial recycling of soil organic matter and pyromineralisation (Findlay et al. 2022), why regularly burned grasslands have sustained for millennia (and continue to maintain) the production of new foliar protein after a burn (Everson and Everson 2016), despite substantial losses of N through recurrent combustion (Hobbs et al. 1991).

Our hypothesis that the *nifH* gene would occur more frequently with the roots of species with a slow, conservative growth strategy that are most tolerant of low soil N (Marques et al. 2017), particularly *A. junciformis*, was not supported as the four study species did not differ in mean gene copy number.Therefore, they all could potentially rely to some extent on ANF in the unfertilised study grassland which has low soil N (∼0.22%; Supplementary Table 1). On richer soils and with addition of mineral fertiliser N, the species with the most acquisitive growth strategy, *E. curvula*, would probably depend more on extant soil N (Hu et al. 2020) than the more metabolically expensive process of BNF (Roley et al. 2018) because of its high physiological capacity to respond to available soil N (Morris 2016). Addition of mineral fertiliser N also reduces diazotroph populations and their colonisation of plant tissues, the types of root exudates most beneficial to them, and their nitrogenase activity, thereby diminishing ANF (Keuter et al. 2014; Marques et al. 2017; Roley et al. 2019; Hu et al. 2020). Careful research is required to examine trade-offs in ANF and competitive ability of mesic grasslands among soil types and along nutrient gradients induced by fertiliser or intensive grazing, as well as the long-term response of ANF to nutrient addition (Zheng et al. 2019).

Another interesting finding was that *nifH* gene copy number was systematically higher (+60%) for all four species in the fresh-compared to the senescent sward (Figure 2). This suggests a positive general effect of fire on potential ANF. There are, however, a few caveats to this result. First, burning regime was confounded with spatial location; however, soil fertility was similar between the two halves of the field (Supplementary Table 1). Second, gene abundance does not always reliably predict in situ N_2_ fixation rates because DNA extractions could contain active and dormant diazotrophs (Wallenstein and Vilgalys 2005) and fixation is strongly mediated by diazotroph community composition and affected by soil environment and resources (esp. N, C, P, Mo), also varying between species and plant parts (Reed et al. 2011; Smercina et al. 2019). *NifH* genes could have been most abundant the season following the fire because fire stimulates N_2_-fixation by rendering phosphorus more available to diazotrophs (Eisele et al. 1989) when the demand for N from rapidly growing grasses would be high, which, in turn, provides carbon-rich root exudates to fuel diazotrophs (Dart 1986). Nitrogenase activity is highest when grasses are actively photosynthesising (Dommergues et al. 1973) and soils are wet (Hu et al. 2020) but Roley et al. (2018) reported peak ANF after senescent plants had translocated nutrients from shoots to roots, providing a potential pool for soil N for spring regrowth. Such seasonal variation in ANF in mesic grasslands in South Africa has yet to be explored.

The diazotrophic bacterial species associated with the four mesic grass species in our study were not identified, but the sequencing of a small sample of amplified gene gene fragments indicated that they were at least of *nifH* gene origin. This gene is relatively conserved, i.e., sequences are similar, ranging across several taxa such as *Azotobacter, Herbaspirillum*, and *Bradyrhizobium* species (Dai et al. 2014). Therefore, due to the short length and level of ambiguities in the sequenced fragments, it was not possible to confidently infer taxonomic assignments from the sequences. However, it is known that strains in the genus *Bradyrhizobium* are not only typical soybean symbionts but also commonly associates with non-legumes and prairie grasses (Bahulikar et al. 2014; Roley et al. 2019; Yoneyama et al. 2019) and the genus is a reliable indicator of grassland diazotrophic communities (Zhu et al. 2022). Although the amplicon sequencing was not taxonomically informative, it confirmed that the *nifH* gene was present and associated with the roots of the study grasses and the differences between sequenced samples revealed several distict *nifH* gene sequences. Further research is required to identify the range of *nifH* gene bearing organisms harbouring these genes on local mesic grasses. A molecular study of 16S rRNA genes (Fonseca-López et al. 2020) could reveal species-rich assemblages of diazotrophs living as endophytes in, or epiphytically on, the roots of local grasses because such communities are typically diverse (Bahulikar et al. 2014; Keymer et al. 2014; Gupta et al. 2019). Diazotroph community composition also varies among plant parts and grass species (Gupta et al. 2014; Wemheuer et al. 2017; Roley et al. 2019). Such variation should be examined in South African grasslands. Also unknown are the effects of plant growth stimulants produced by diazotrophs (Dobbelaere et al. 2003; Smercina et al. 2019) and their potential synergism with mycorrhiza on the N and P nutrition and productivity of mesic grasses (Bauer et al. 2012)

Grassland ANF could become more critical with continued oligotrophication of ecosystems resulting from insufficient soil N to meet the higher demands of faster-growing plants fuelled by steadily rising levels of atmospheric CO_2_ (Craine et al. 2018). Carbon supply and temperature constraints to ANF could be respectively alleviated by warming and elevated [CO_2_] but drying with climate change would retard BNF because of the strong control that soil moisture exerts on N_2_-fixation rates (Reed et al. 2011; Khan et al. 2021). Therefore, the future dynamics of ANF dynamics in grassland and consequences for N availability to grasses (Smercina et al. 2019) are uncertain.

Furthermore, mesic grasslands could be sources of culturable nitrogen fixing bacteria that could be utilised for enhancing nitrogen supply in cereal crops (Ke et al, 2019; Bennett et al. 2020) and cultivated pastures used for animal production (Reis et al. 2001), thereby reducing requirements for expensive synthetic nitrogen fertilizers which contribute to CO_2_ emissions (Chai et al, 2019).

## Conclusion

This study provides the first genetic evidence that bacterial diazotrophs could commonly occur in South African mesic grassland and that these N_2_-fixing microbes can associate with the roots of a wide range of grass species differing in their nitrogen economies. Therefore, ANF could play an underappreciated significant role in grassland ecosystems. The findings also open an interesting and important research agenda to understand what diazotroph species are involved in grass ANF, where they occur on or inside grasses, how much nitrogen they fix and supply to grasses and to the whole grassland ecosystem, how ANF is affected by soil nutrient status and common perturbations such as drought, grazing, fire, and mowing, and how diazotrophic communities could be altered by climate change.

## Acknowledgements

We are grateful to the following people who assisted with the collection, preparation, and preliminary analysis of samples: Naledi Zama, Tinta Morris, Heather Tredgold, Matthew van Wyngaard, Steven O’Connor, and Michael Relihan. We thank the Analytical Services Sub-directorate of the KwaZulu-Natal Department of Agriculture and Rural Development for undertaking the soil analysis.

## Additional material

Supplementary Table 1 – Soil analysis

## Notes

### Competing Interest Statement

The authors have declared no competing interest.

