## Supplementary Table 1 for "Associative nitrogen fixation could be common in South African mesic grassland"

**Supplementary Table 1:** Soil chemical characteristics of composite samples (n = 10) of topsoil (150 mm depth) from the 'Old' and 'New' positions of the study site. Soil analyses were undertaken by the Analytical Services Sub-directorate of the KwaZulu-Natal Department of Agriculture and Rural Development, Cedara, Hilton, South Africa (Manson et al. 2020).

| POSITION | SAMPLE DENSITY | P | K | Ca | Mg | Total cations | Exch. acidity | Acid sat. | pH | Zn | Mn | Cu | Carbon | Nitrogen | C/N |
|---|---|---|---|---|---|---|---|---|---|---|---|---|---|---|---|
| | | mg L-1 | | | | | | % | | mg L-1 | | | % | % | |
| **OLD** | 0.94 | 6 | 165 | 1109 | 504 | 10.25 | 0,15 | 1 | 4,82 | 3,3 | 57 | 5,3 | 3,73 | 0,23 | 16,3 |
| **NEW** | 1.04 | 8 | 225 | 1033 | 430 | 9.41 | 0,15 | 1 | 5,03 | 4,7 | 46 | 6,4 | 3,29 | 0,21 | 15,8 |

### Reference


Manson AD, Bainbridge SH, Thibaud GR. Methods used for analysis of soils and plant material by Analytical Services at Cedara. South Africa: KwaZulu-Natal Department of Agriculture and Rural Development, South Africa..
